# Identification of sex differences in tumor-specific T cell infiltration in bladder tumor-bearing mice treated with BCG immunotherapy

**DOI:** 10.1101/2020.06.19.161554

**Authors:** Matthieu Rousseau, Conan J O O’Brien, Eduardo Antequera, Hana Zdimerova, Dilay Cansever, Tracy Canton, Anna Zychlinsky Scharff, Molly A Ingersoll

**Affiliations:** Department of Immunology, Institut Pasteur, 75015 Paris, France; INSERM U1223, 75015 Paris, France

## Abstract

Bladder cancer is the fourth most common cancer for men. However, women are often diagnosed with later stage disease and have poorer outcomes. Whether immune-based sex differences contribute to this discrepancy is unclear. In addition, models to investigate tumor-specific immunity in bladder cancer, in the context of tumor development or response to therapy, are lacking. To address this specific unmet need, we incorporated a commonly used model antigen, ovalbumin, into two well-established models of bladder cancer; the orthotopic MB49 cell line model and the carcinogenic BBN bladder cancer model. We tested the utility of these models to investigate tumor-specific immunity in the context of immunotherapy in both sexes. We found that BCG vaccination, prior to weekly BCG instillation does not impart an immune-specific benefit to tumor-bearing mice in the context of multiple BCG instillations. Furthermore, tumors developed in the testes in male mice, precluding the use of the MB49 model to directly investigate sex-based immune differences. In the BBN model, we observed that more tumor antigen-specific CD8^+^ T cells infiltrated male bladders compared to female bladders in the context of BCG immunotherapy and that these cells had the highest levels of the exhaustion marker PD-1. We propose our modified BBN model will contribute to our understanding of how tumor-specific immunity arises in bladder cancer. Additionally, the BBN bladder cancer model may help to uncover sex differences in tumor-specific immunity, which would provide valuable information for the development of new treatments or combination therapies for bladder cancer in women and men.

## Introduction

Bladder cancer is an exceedingly common disease comprised of diverse clinical and molecular subgroups that is approximately three times more prevalent in men than women (Felsenstein and Theodorescu, 2018). At initial diagnosis, ~75% of cancers are nonmuscle invasive with different grades, stages and risk levels (Sanli et al., 2017). Nonmuscle invasive bladder cancer (NMIBC) has highly variable five year recurrence and progression rates following transurethral tumor resection, however these rates are mitigated by Bacillus Calmette-Guérin (BCG) immunotherapy (Brandau and Suttmann, 2007; Pettenati and Ingersoll, 2018). Intravesical administration of BCG, a passage-attenuated strain of *Mycobacterium bovis*, has been the standard of care for NMIBC since Morales and colleagues demonstrated in 1976 that it reduces tumor recurrence (Morales et al., 1976; Pettenati and Ingersoll, 2018). Subsequent clinical studies show that patients with NMIBC have better clinical outcomes and progression-free survival when BCG immunotherapy follows transurethral tumor resection (Brandau and Suttmann, 2007; Pettenati and Ingersoll, 2018).

Although BCG immunotherapy has been used to treat NMIBC for more than 40 years, the mechanism by which it mediates tumor immunity is still incompletely understood (Kamat et al., 2018; Pettenati and Ingersoll, 2018). It is clear, however, that BCG induces a complicated interplay of innate and adaptive immune responses and cross-talk among many cell types. Lasting protection requires CD4^+^ and CD8^+^ T cells, as the absence of these populations, either individually or together, abolishes BCG-mediated anti-tumor immunity (Ratliff et al., 1987; Ratliff et al., 1993). These effector T cell responses are shaped by innate immunity. For example, neutrophils, which mediate acute inflammation, induce T cell migration, and neutrophil depletion significantly impairs CD4^+^ T cell trafficking to the bladder, abrogating the antitumor activity of BCG (Suttmann et al., 2006). Infiltrating monocyte-derived cells encounter a suppressive tumor microenvironment that induces their polarization to alternatively-activated macrophages, which may favor tumor development (Mariathasan et al., 2018; Sjodahl et al., 2014). These M2-like cells contribute to an immunosuppressive type 2 immune response, which in turn shapes T helper (T_h_) cell bias. This is particularly relevant in bladder cancer, as the T_h_1:T_h_2 ratio, and more specifically, the bias of CD4^+^ T cells towards a T_h_1 profile, is crucial for BCG-mediated tumor immunity. Indeed, BCG immunotherapy is ineffective in mice deficient for the T_h_1 cytokines IFNγ and IL-12, but is improved in mice lacking the Th2 cytokine IL-10 (Riemensberger et al., 2002).

BCG immunotherapy induces long lasting immunity, however the antigens recognized by the immune system, particularly in early stages, are largely unknown. BCG immunotherapy induces immunity to the bacteria themselves, which may contribute to its efficacy, patient seroconversion during BCG therapy is correlated with better outcomes (Kelley et al., 1985; Lamm et al., 1981). Pre-existing immunity to BCG from childhood vaccination may also enhance response to therapy (Biot et al., 2012). This can be modeled in animals by vaccination prior to tumor induction. For example, compared to control-treated animals, rats pre-sensitized to BCG have reduced tumor progression after BCG intralesional injection. BCG-vaccinated dogs have a greater inflammatory response including increased immune cell infiltration into the bladder following BCG instillation (Bloomberg et al., 1975; Lamm et al., 1977). In mice, BCG vaccination induces accelerated T cell infiltration into the bladder following a single intravesical instillation of BCG and; confers a survival advantage to vaccinated mice over nonvaccinated mice in an orthotopic tumor model (Biot et al., 2012). Humans with pre-existing BCG immunity have improved outcomes, such as longer recurrence-free survival intervals, following immunotherapy (Biot et al., 2012; Lamm et al., 1981). BCG immunity is, however, an imperfect surrogate for tumor immunity in preclinical models and in patients.

As additional immunotherapeutic approaches, including checkpoint inhibition, demonstrate some success in the treatment of bladder cancer (Ghatalia et al., 2018), it is vital to understand how tumor antigen-specific immunity is influenced by immunotherapy, particularly to dissect why these therapies are not always successful (Philip et al., 2020). Critically, models and tools to investigate tumor-specific immunity in bladder cancer are lacking. Our goal for this study was to develop an optimized preclinical bladder cancer model in which we could assess the influence of immunotherapy on tumor-specific immunity in both sexes. As bladder cancer is approximately three times more common in men than in women, and innate bladder mucosal responses differ between the sexes in infection, we used female and male mice in both models to uncover potential sex-based differences in tumor immunity (Antoni et al., 2017; Zychlinsky Scharff et al., 2019). To develop these models, we incorporated a known model antigen into two well-established models of bladder cancer. We measured immune responses, including tumor-antigen specific responses, following BCG and checkpoint inhibition in female and male mice. We found that immunity differs depending upon the model used, and that immunotherapeutic approaches may impact the adaptive immune responses differently in male and female mice.

## Results

### BCG Connaught and BCG Danish induce robust T cell priming in vivo

Previously, we demonstrated that the BCG Connaught substrain induces more robust CD8^+^ T cell priming in C57BL/6 mice compared to BCG Tice, and correlates with superior protection in bladder cancer patients (Rentsch et al., 2014). To determine whether other substrains induce variable T cell priming, we vaccinated 6 week old female C57BL/6 mice subcutaneously with PBS or BCG strains Connaught, Tice, Danish, China, Japan, or RIVM (Medac). We analyzed splenocytes after 15 days using D^b^-Mtb32 tetramers, which bind the TCR of CD8^+^ T cells specific for the *Mycobacterium* epitope GAPINSATAM (Irwin et al., 2005). BCG Connaught and Danish substrains were the only strains to significantly induce BCG-specific CD8^+^ T cell priming over PBS-treated mice (**Figure 1**). BCG substrains Tice, China, Japan, and RIVM did not induce the development of BCG-specific T cells over the proportion or number of specific T cells observed in PBS-treated animals (**Figure 1**). Notably, BCG Connaught induced the greatest number of BCG tetramer-positive T cells, and as such, we chose to use this strain to investigate the induction of BCG-mediated tumor immunity in this study.

**Figure 1.**
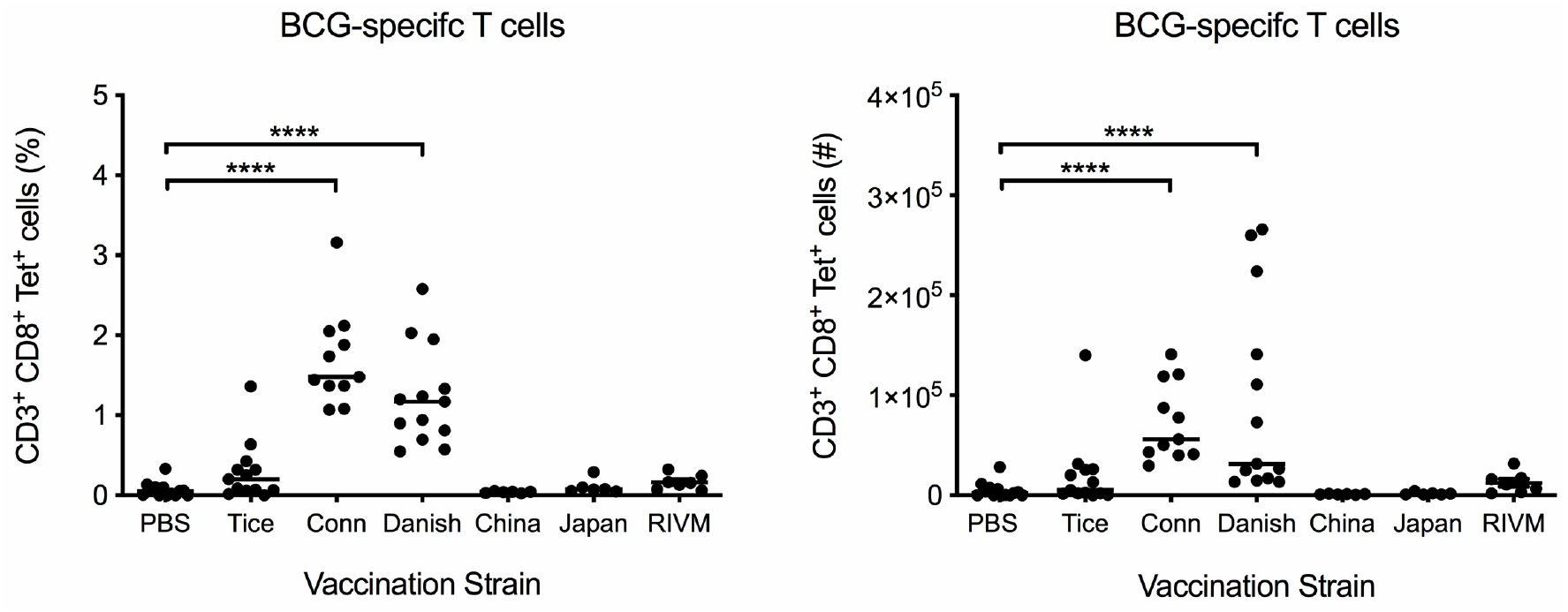
BCG Connaught and Danish induce robust T cell priming in vaccinated mice. Six week old female C57BL/6 mice were vaccinated subcutaneously with PBS or 3×10^6^ CFU of BCG substrains Tice, Connaught, Danish, China, Japan or RIVM (Medac). BCG-specific CD8^+^ splenic T cells were identified by flow cytometry using D^b^-Mtb32 tetramers 15 days postvaccination. Graphs depict the percentage and number of BCG-specific CD8^+^ T cells in mice vaccinated with each strain. Each dot represents one mouse, lines are medians. Data are pooled from three independent experiments. ****p<0.0001, Kruskal-Wallis test comparing each strain to the PBS control group with Dunn’s post-test to correct for multiple comparisons.

### BCG vaccination does not induce BCG-specific T cell infiltration into bladders

In BCG-vaccinated mice, T cell infiltration into the bladder following a single intravesical instillation of BCG is equivalent to the level of T cell infiltration observed in unvaccinated mice instilled three times with BCG (Biot et al., 2012). As patient populations, naïve or not to BCG, receive more than one intravesical BCG instillation in the context of an induction cycle, we tested whether multiple BCG instillations would induce even greater T cell infiltration in vaccinated animals. We subcutaneously injected 6 week old female C57BL/6 mice with PBS or BCG and, 2-3 weeks later, treated all animals with intravesical BCG once per week for three weeks (**Figure 2A**). We analyzed bladder-associated antigen-presenting and effector cell populations by flow cytometry 24 hours after the final instillation. BCG vaccination did not alter the total cellularity or the number of CD45^+^ immune cells in the bladder (**Figure 2B**). Resident macrophage and dendritic cell (DC) populations were not different between PBS and BCG-vaccinated mice treated three times with intravesical BCG (**Figure 2B**). Vaccinated animals had a significant increase in the number of mature (*i.e.* MHC II^+^) monocyte-derived cells recruited from circulation compared to PBS-treated mice (**Figure 2C**). Finally, while there were no differences in total CD3^+^ T cell recruitment, CD4^+^ T cells were significantly increased in BCG-vaccinated mice compared to the PBS-treated group (**Figure 2C**).

**Figure 2.**
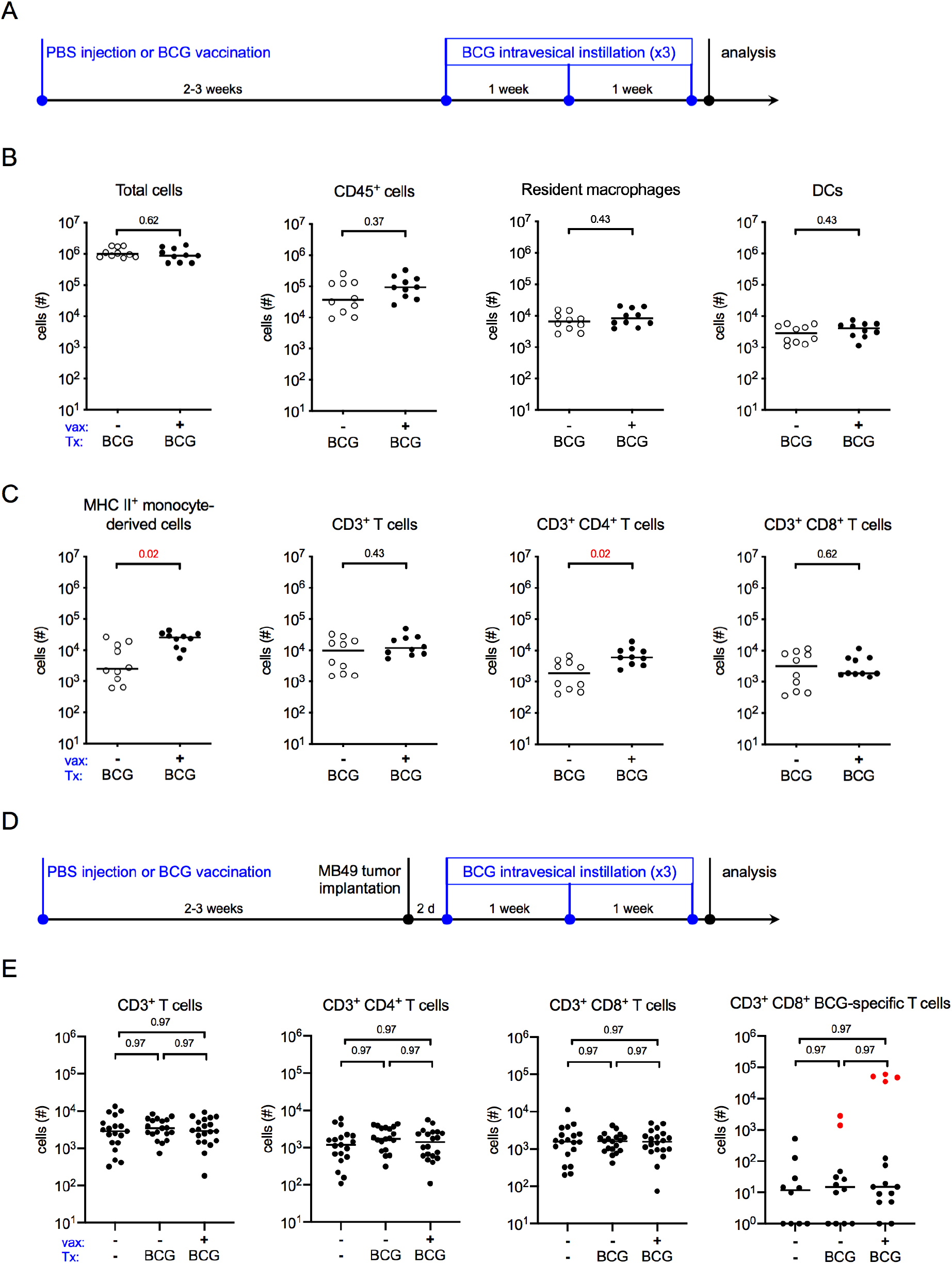
BCG-specific T cells do not infiltrate the bladder. (A) Schematic of the experiment depicted in **B** and **C**. Briefly, six week old female C57BL/6 mice were vaccinated subcutaneously with PBS or 3×10^6^ CFU of BCG Connaught. Two to three weeks later the mice received 5×10^6^ CFU of BCG Connaught intravesically once a week for three weeks. 24 hours after the last instillation, the bladders are analyzed by flow cytometry. (**B** and **C**) Graphs depict total specified immune cell populations. (**D**) Schematic of the experiment depicted in **E**, which is the same as shown in **A**, except two to three weeks post-vaccination mice received 2×10^5^ MB49 cells by intravesical instillation. PBS or BCG Connaught administration was started two days after tumor implantation. One group received no injections or intravesical instillations. 24 hours after the last instillation, the bladders are analyzed by flow cytometry. (**E**) Graphs depict total specified immune cell populations. Red dots denote the total number of BCG-specific CD8^+^ T cells found in the spleens of vaccinated mice. Data are pooled from 2-3 experiments, *n*=5-7 mice per experiment. Each dot represents 1 mouse, lines are medians. Significance was determined using the nonparametric Mann-Whitney test to compare vaccinated to unvaccinated mice for each immune population shown, and the p-values were corrected for multiple testing using the false discovery rate (FDR) method. All calculated/corrected p-values are shown and those meeting the criteria for statistical significance (p<0.05) are depicted in red.

We next considered whether vaccination would also improve effector cell infiltration in the context of a tumor. Six week old female C57BL/6 mice were subcutaneously injected with PBS or BCG and 2-3 weeks later, orthotopically implanted with the syngeneic tumor cell line MB49, which expresses luciferase. Tumor implantation was verified by bioluminescent imaging at day 7 post-implantation. PBS control-treated and BCG-vaccinated mice were treated intravesically with BCG 2 days after tumor implantation, and then once per week for a total of three treatments (**Figure 2D**). A third group, which was injected with PBS at the start of the experiment, was implanted with MB49 but not treated with intravesical therapy. Similar to tumor-naïve mice in **Figure 1C**, we were unable to identify differences in total CD3^+^ T cell infiltration between tumor-bearing PBS-treated and BCG-vaccinated mice receiving BCG intravesical therapy, and, surprisingly, we observed no differences between these groups and tumor-bearing animals receiving no treatment at all (**Figure 2E**). In addition, there were no differences in CD4^+^ or CD8^+^ T cell infiltration among the three groups, suggesting that in this model, effector cell infiltration is tumor-driven, rather than induced by BCG (**Figure 2E**). We hypothesized that although total T cell numbers were not different, BCG-specific T cell infiltration may differ among the groups. Using D^b^-Mtb32 tetramers, we identified a very small population of BCG-specific CD8^+^ T cells in bladder tissue that was not different among the groups. Furthermore, the number of tetramer^+^ CD8^+^ T cells in bladders was more than 1000 times lower than the number of tetramer^+^ T cells in the spleens of vaccinated animals (**Figure 2E**, compare red dots to black). This finding suggests that T cells in the bladder are not BCG-specific T cells. The specificity of these T cells, in particular towards tumor antigens however, is unclear.

### A model antigen-containing MB49 cell line, MB49^mCh-ova^, induces immune cell infiltration

One limitation of the MB49 orthotopic tumor model is that it cannot be used to directly investigate tumor-specific immunity because antigens arising from the tumor are not known. We transduced an MB49 cell line to express mCherry red fluorescent protein (mCh) fused to ovalbumin (ova), a widely-used model antigen, which can be followed using transgenic mouse tools (Barnden et al., 1998; Hogquist et al., 1994). The resultant strain, MB49^mCh-ova^, also expressed luciferase, permitting tumor detection in live mice by *in vivo* imaging (**Figure 3A**). Red fluorescent protein was easily detected by flow cytometry (**Figure 3B**). To test whether modification of MB49 altered its ability to induce inflammation or form tumors, we assessed tumor implantation and immune cell infiltration in mice using either the parental MB49 strain or MB49^mCh-ova^. We calculated the implantation rates (percentage of success) by dividing the number of mice with tumors by the total number of mice implanted per experiment, and while implantation was more variable in MB49^mCh-ova^-implanted mice compared to mice receiving MB49, this likely reflects investigator experience as implantation rates of MB49^mCh-ova^ improved over time (**Figure 3C**, the order of experiments is shown). Importantly, in all analyses, we only included animals with tumors verified by *in vivo* imaging at 7 days post-tumor instillation, because it cannot be determined whether an immune response eradicated the small tumor burden prior to day 7 or tumor implantation failed in these animals. With respect to immune cell populations, we observed no differences between MB49 and MB49^mCh-ova^-implanted mice in total CD45^+^ cells; resident macrophage and DC populations; infiltrating myeloid cells, such as mature (MHC II^+^) and immature (MHC II^-^) monocyte-derived cells, eosinophils, or neutrophils; or T lymphocytes (**Figure 3D**), indicating the mCherry-ova transgene did not impact MB49-induced immune cell infiltration.

**Figure 3.**
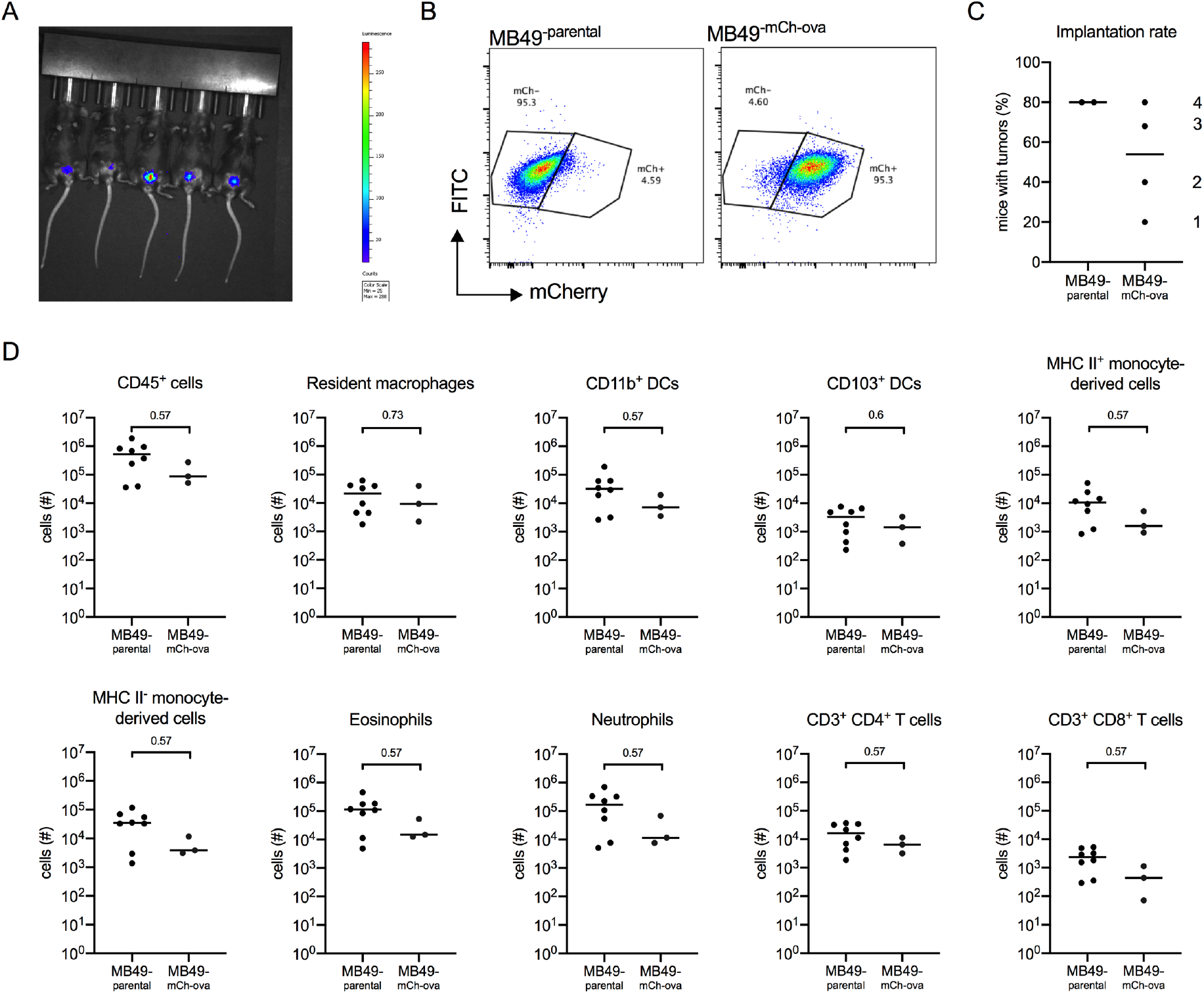
The MB49^mCh-ova^ cell line induces immune cell infiltration. Six week old female C57BL/6 mice were instilled with 2×10^5^ MB49 or MB49^mCh-ova^ cells. Presence of a tumor was verified by *in vivo* imaging and immune cell populations were quantified by flow cytometry on day 14. (**A**) Representative image of tumor detection *in vivo* using the IVIS Spectrum In Vivo Imaging System on day 7 post tumor instillation. (**B**) Dot plots depict mCherry fluorescent protein expression in the parental MB49 cell line and the MB49^mCh-ova^ cell line. (**C**) Graph shows the implantation rate for MB49 or the MB49^mCh-ova^ on day 7 post tumor instillation in C57BL/6 female mice. The numbers indicate the order in which the experiments were performed. (**D**) Graphs depict total specified immune cell populations. Graphs are pooled from 2 experiments, *n*=5 mice per experiment, animals without tumors by day 7 were excluded from analysis. Each dot represents 1 mouse, lines are medians. In (**D**), significance was determined using the nonparametric Mann-Whitney test to compare immune cell populations between the parental and the modified MB49 implanted mice, and p-values were corrected for multiple testing over all analyses using the FDR method.

### BCG vaccination increases immune cell infiltration in combination with checkpoint inhibition

As we did not observe differences in effector cell infiltration in the presence or absence of BCG vaccination in tumor-bearing mice (**Figure 2E**), we tested whether vaccination would alter immune cell infiltration in combination with checkpoint inhibition. Six week old female C57BL/6 mice were vaccinated or not with BCG Connaught, and 2-3 weeks later MB49^mCh-ova^ cells were instilled into the bladder. Two days post-tumor implantation, all mice received BCG intravesically and day 3 post-tumor implantation, all mice received anti-programmed death-ligand 1 (α-PD-L1) antibody intraperitoneally, similar to the dosing schedule in humans (**Figure 4A**). As above, mice without luminescent tumors on day 7 were excluded. BCG intravesical therapy was continued on days 9 and 16 post-tumor implantation and α-PD-L1 antibody was given on days 6 and 9 post-tumor implantation for a total of three treatments with each therapy (**Figure 4A**). We analyzed bladders 24 hours after the final BCG treatment by flow cytometry. There were no differences in the number of CD45^+^ immune cells between vaccinated and unvaccinated mice (**Figure 4B**). We also observed no changes in the quantity of cells with antigen-presenting capabilities, such as resident macrophages, DC subsets, or mature MHC II^+^ monocyte-derived cells between unvaccinated and vaccinated mice (**Figure 4C**). Infiltrating CD4^+^ T cell numbers were not different between the two groups, however there were significantly more CD8^+^ T cells in the vaccinated group (**Figure 4D**). Surprisingly, we found that in the context of intravesical BCG and α-PD-L1 antibody treatment, vaccinated mice had greater myeloid cell infiltration, including immature MHC II^-^ monocyte-derived cells, eosinophils, and neutrophils, into the bladder compared to unvaccinated mice (**Figure 4E**), suggesting that vaccinated mice have a more pro-inflammatory bladder microenvironment.

**Figure 4.**
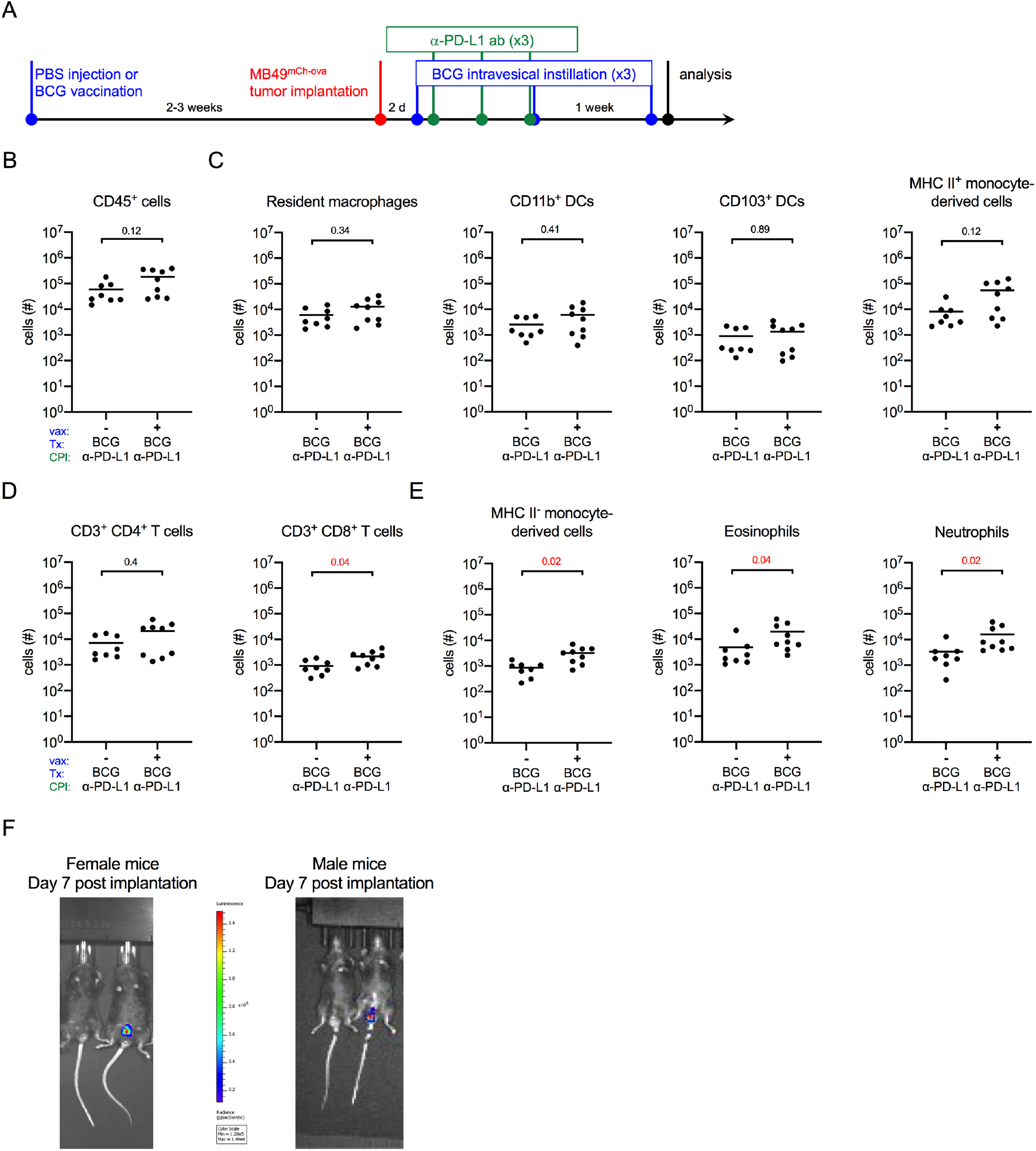
BCG vaccination combined with checkpoint immunotherapy increases innate immune cell, and CD8^+^ T cell infiltration. (**A**) Schematic of the experiment depicted in **B-E**. Six week old female C57BL/6 mice were vaccinated with PBS or 3×10^6^ CFU of BCG Connaught and instilled with 2×10^5^ MB49^mCh-ova^ cells 2-3 weeks after. All mice received 5×10^6^ CFU of BCG Connaught intravesically two days post-tumor instillation and then once a week for a total of 3 instillations. On day 3, 6, and 9 post tumor instillation, all the mice received α-PD-L1 immunotherapy (indicated as CPI in the X-axes) by intraperitoneal injection. 24 hours after the last BCG instillation, bladder immune cell populations were analyzed by flow cytometry. (**B**, **C**, **D**, and **E**) Graphs depict total specified immune cell populations. (**F**) Representative image of *in vivo* tumor detection in female and male mice 7 days post-tumor implantation. In **B**, **C, D,** and **E,** data are pooled from 2 experiments, *n*=4-5 mice per experiment, animals without tumors by day 7 were excluded from analysis. Each dot represents 1 mouse, lines are medians. Significance was determined using the nonparametric Mann-Whitney test to compare vaccinated to unvaccinated mice for each immune population shown, and the p-values were corrected for multiple testing using the FDR method. All calculated/corrected p-values are shown and those meeting the criteria for statistical significance (p<0.05) are in red.

Typically, only female mice are used for the MB49 mouse model as catheterization of male mice is reported to be unattainable by many preclinical bladder studies (El Behi et al., 2013; Hagberg et al., 1983; Oliveira et al., 2009; Olson et al., 2016; Seager et al., 2009). Given that more men than women develop bladder cancer, and that sex differences contribute to this bias in incidence (Kaneko and Li, 2018), we used a male mouse intravesical instillation protocol that we previously optimized to measure sex-specific differences in innate immunity to urinary tract infection (Zychlinsky Scharff et al., 2017; Zychlinsky Scharff et al., 2019). We implanted female and male mice with MB49 to determine whether sex differences impact immunity or response to BCG. We monitored tumor growth by *in vivo* luminescence imaging. To our great disappointment, while tumors were detected in the bladders of female mice, in male mice, luminescence was detected exclusively in the testes (**Figure 4F**). In summary, our results suggest that the MB49 model is inappropriate to address sex differences, and that while it may be that BCG vaccination enhances BCG immunotherapy in combination with checkpoint inhibition, it is difficult to easily assess these subtle differences as the rapid tumor growth of MB49 or MB49^mCh-ova^ renders this orthotopic model of limited utility.

### The BBN model permits study of sex-specific immune responses to immunotherapy

As we eliminated the MB49^mCh-ova^ model as a means to investigate potential sex-specific differences in response to immunotherapy, we turned to the N-butyl-N-(4-hydroxybutyl) nitrosamine (BBN) bladder cancer model. BBN, a carcinogen that specifically promotes the development of bladder cancer, has been used to investigate tumor development for more than 50 years (Bertram and Craig, 1972; Druckrey et al., 1964). Notably, this model has not been extensively exploited for investigation of the immune response to tumor development. To follow tumor-specific immunity, we used the transgenic URO-OVA mouse, which expresses ovalbumin from the bladder-specific uroplakin II promoter (Liu et al., 2007). Importantly, in the absence of activated transgenic CD8^+^ OT1 T cells, which recognize ovalbumin, the UROOVA mouse develops normally (Liu et al., 2007). We reasoned that BBN-treated URO-OVA mice would develop tumors expressing an endogenous “tumor” antigen, ovalbumin. By adoptively transferring CD8^+^ OT1 T cells, we could follow development of tumor-specific immunity in the context of immunotherapy (**Figure 5A**). Additionally, this model can be used to investigate sex differences. To validate this approach, we recapitulated the finding that BBN treatment induces larger tumors in male mice compared to female mice given BBN for the same period of time. (**Figure 5B**) (Bertram and Craig, 1972; Kaneko and Li, 2018).

**Figure 5.**
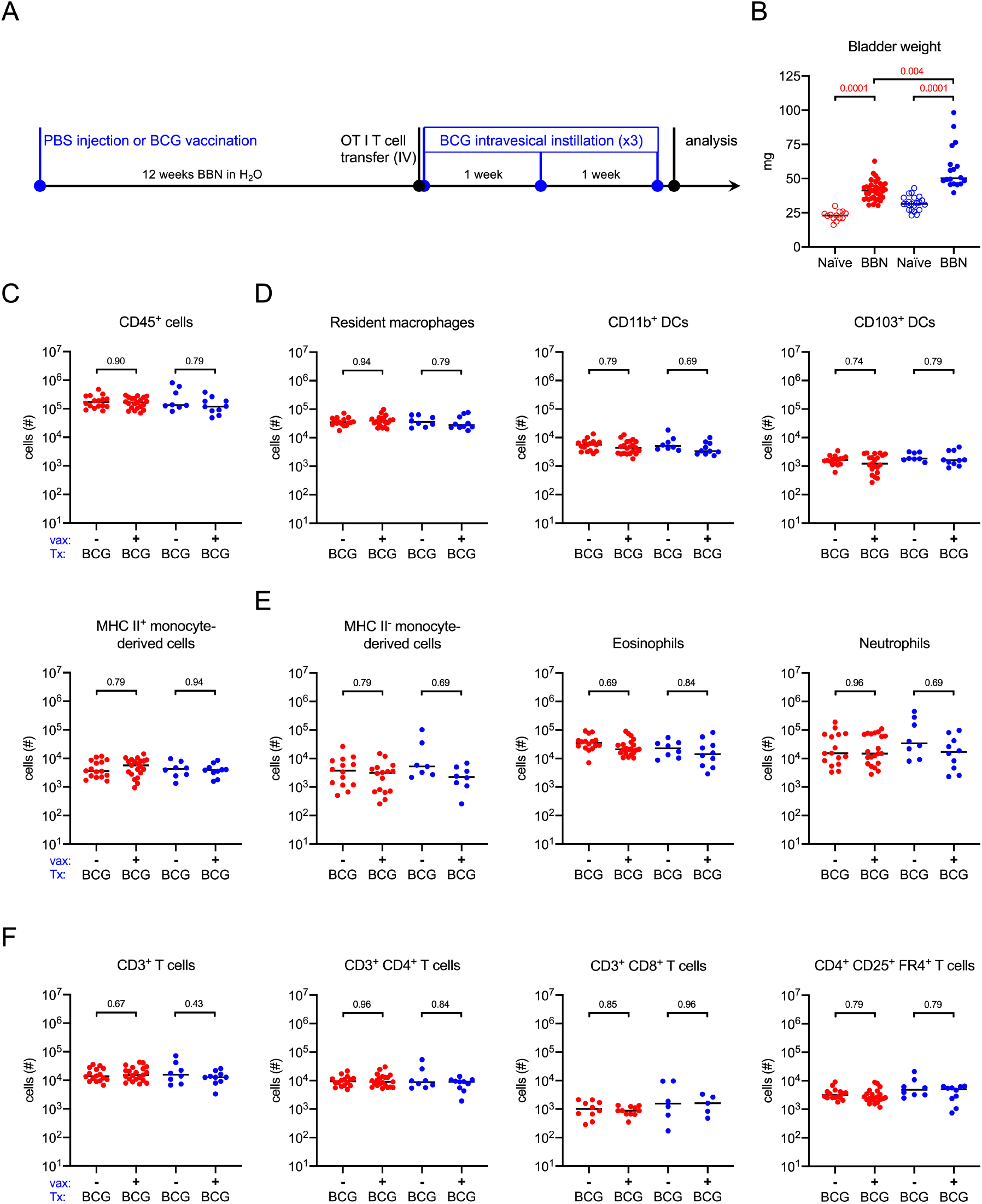
BCG vaccination does not impact immune cell infiltration in the BBN model. (**A**) Schematic of the experiments for data shown in this figure, **Figure 6**, and **Figure 7**. Six week old female (red dots) and male (blue dots) URO-OVA mice were injected subcutaneously with PBS or 3×10^6^ CFU of BCG Connaught and given 0.05% BBN in the drinking water for the duration of the experiment. OT1 splenocytes were transferred by intravenous injection at week 12 and mice were treated with 5×10^6^ CFU of BCG Connaught intravesically once a week for three weeks. 24 hours after the last BCG instillation bladder immune cell populations were analyzed by flow cytometry. (**B**) Graph displays the bladder weight of naïve or tumor bearing female and male URO-OVA mice. (**C**, **D**, **E**, and **F**) Graphs depict total specified immune cell populations. Graphs are pooled from 4 experiments, *n*=1-6 mice per experiment. Each dot represents 1 mouse, lines are medians. Significance was determined using the nonparametric Kruskal-Wallis test to compare vaccinated to unvaccinated mice for each immune population shown, and the p-values were corrected for multiple testing using the FDR method. All calculated/corrected p-values are shown and those meeting the criteria for statistical significance (q<0.05) are depicted in red.

### Immune cell infiltration is not altered in BCG-vaccinated BBN tumor-bearing URO-OVA mice

Having established that BBN induces tumor growth in the URO-OVA mouse, we tested whether immunity to BCG would impact immune cell infiltration in the BBN model, as, to the best of our knowledge, this has not been previously reported. Six week old female and male URO-OVA mice were vaccinated or not with BCG and given 0.05% BBN in the drinking water for 12 weeks. At week 12, 1×10^6^ CD8^+^ OT1 transgenic T cells were adoptively transferred into all mice and at 12, 13, and 14 weeks, mice were treated with BCG intravesically (**Figure 5A**). This timepoint was chosen to favor smaller tumors that are not yet muscle-invasive (Bertram and Craig, 1972; Druckrey et al., 1964). Mice were sacrificed for analysis by flow cytometry 24 hours after the final BCG instillation. Surprisingly, we detected no statistically significant differences between unvaccinated and vaccinated mice of either sex in the number of CD45^+^ immune cells (**Figure 5C**). We also observed no changes in antigen-presenting cells (APCs), including resident macrophages, DC subsets, or mature MHC II^+^ monocyte-derived cells between unvaccinated and vaccinated mice (**Figure 5D**). The number of infiltrating immature MHC II^-^ monocyte-derived cells, eosinophils, and neutrophils was not different between the two groups in either sex (**Figure 5E**). Finally, there were no differences in total T cells, or CD4^+^ and CD8^+^ T cell subsets (**Figure 5F**). In these analyses, we included an antibody against folate receptor 4 (FR4), which is highly expressed on natural regulatory T cells (Yamaguchi et al., 2007), and while we detected a large number of CD4^+^ regulatory T cells, there were no differences between unvaccinated and vaccinated mice in either sex (**Figure 5F**). As statistical analysis supported that these data did not arise from different distributions (range of p values=0.43-0.96 for all populations measured) in any immune cell populations investigated, we combined the data from unvaccinated and vaccinated animals for further analysis.

### CD4^+^ and CD8^+^ T cells upregulate PD-1 in tumor bearing mice

Checkpoint inhibition immunotherapy in bladder cancer has already shown great potential, (Powles et al., 2014; Rosenberg et al., 2016; Sharma et al., 2016). As checkpoint molecules are upregulated during immune responses, particularly in chronic inflammation and during checkpoint inhibition immunotherapy in humans (Pardoll, 2012), we first compared the expression of PD-L1 on innate immune cells and PD-1 on effector T cells from the mice in **Figure 5** to naïve age and sex matched animals. We observed that PD-L1 was increased on neutrophils, but not eosinophils or immature monocytes, in BBN-treated mice compared to naïve mice in both sexes (**Figure 6A**). Among APCs, cells likely to engage effector T cells in the tumor environment, PD-L1 expression was not significantly elevated on mature MHC II^+^ monocyte-derived cells, however, resident macrophages and both subsets of DCs in BBN-treated mice had increased PD-L1 levels compared to naïve animals of both sexes (**Figure 6B**). In addition, we observed that PD-1 expression was elevated on CD4^+^ and CD8^+^ T cells in both sexes of mice exposed to BBN compared to naïve mice (**Figure 6C**). We were surprised to observe PD-1 expression was not higher on regulatory T cells in BBN-treated mice compared to naïve animals (**Figure 6C**). Finally, OT1 T cells expressed the highest levels of PD-1 of all T cell populations measured (**Figure 6D**). As these cells were handled *ex vivo*, we questioned whether these cells had upregulated PD-1 prior to adoptive transfer. We measured PD-1 expression on CD8^+^ OT1 T cells just prior to transfer, observing that their PD-1 expression was lower than that observed on CD8^+^ OT1 T cells in BBN-treated mouse bladders (**Figure 6D**). Finally, although these checkpoint molecules were significantly increased on a number of immune cell populations, no differences were observed in their expression between female and male mice exposed to BBN (**Figure 6A-D**).

**Figure 6.**
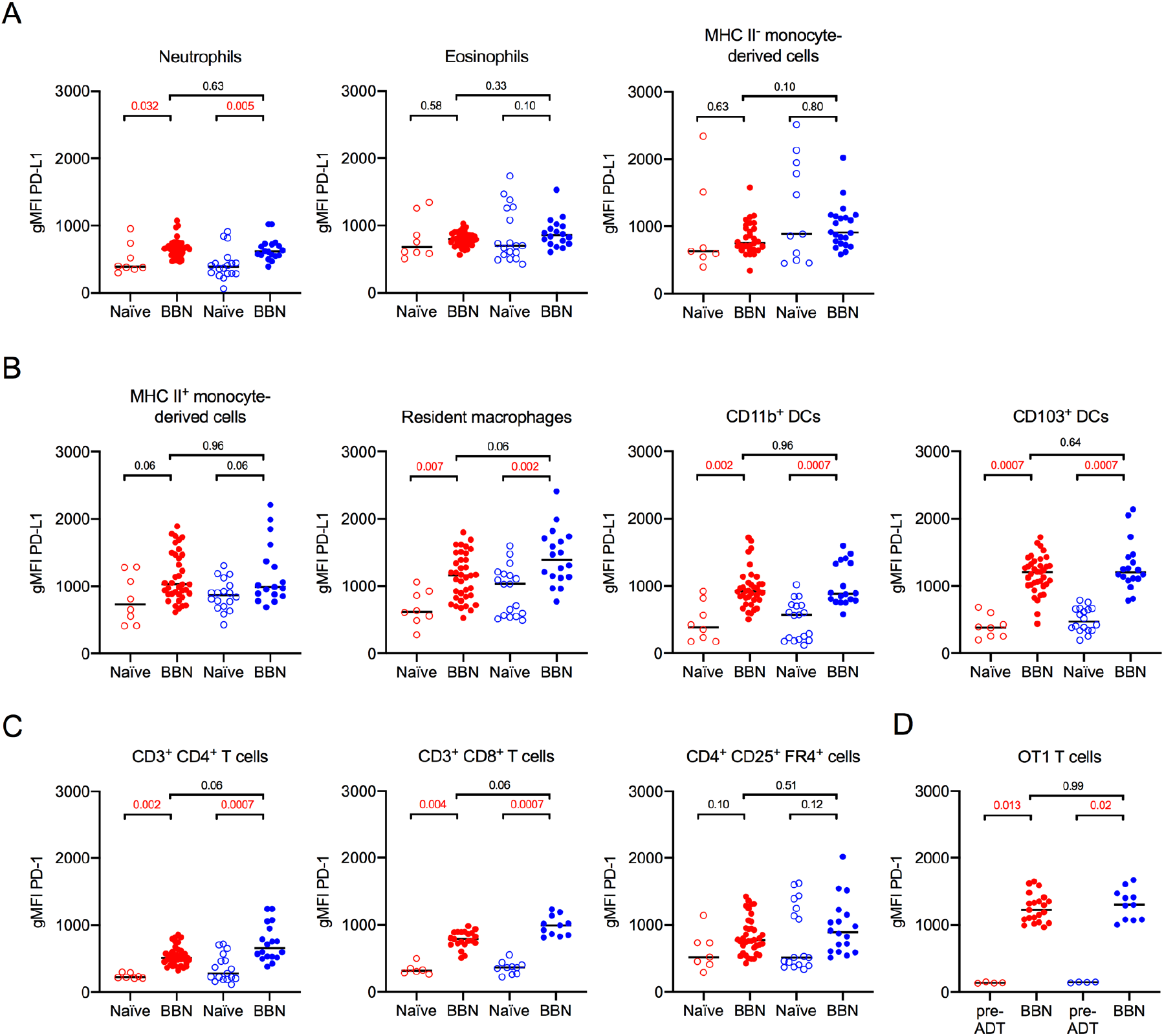
PD-L1 and PD-1 expression increases in male and female tumor-bearing mice. Mice were treated as shown in Figure 5A. (**A**, **B**) Graphs depict the geometric mean fluorescent intensity (gMFI) of PD-L1 or (**C**) PD-1 on the surface of the specified immune cell populations in naïve or tumor bearing female (red dots) and male (blue dots) URO-OVA mice. Graphs are pooled from 10 experiments, *n*=1-7 mice per experiment. Each dot represents 1 mouse, lines are medians. In **6D**, “pre-ADT” = pre-adoptive transfer. Significance was determined using the nonparametric Kruskal-Wallis test to compare naïve to BBN-treated mice for each immune cell population and to compare female to male BBN-treated mice. p-values were corrected for multiple testing using the FDR method. All calculated/corrected p-values are shown and those meeting the criteria for statistical significance (p<0.05) are in red.

### More tumor antigen-specific T cells infiltrate male bladders than female bladders

Finally, we analyzed immune cell infiltration and potential sex differences between female and male mice given BBN for 12 weeks and treated with BCG immunotherapy. Naïve age- and sex-matched animals were included to determine the magnitude of the immune cell infiltrate. Compared to naïve animals, statistically significant CD45^+^ immune cell infiltration was readily apparent in both female and male BBN-treated mice (**Figure 7A**). Innate immune cells, including neutrophils, eosinophils, and immature MHC II^-^ monocyte-derived cells were all increased 10-100 times over naïve mice in both sexes (**Figure 7B**). Interestingly, there were no differences in infiltration of these populations between the sexes, in contrast to what we reported in the context of urinary tract infection, in which female mice have a significantly greater number of immune cells present in their bladders during infection (Zychlinsky Scharff et al., 2019). APCs, including MHC II^+^ monocyte-derived cells, resident macrophages, and both subsets of DCs were significantly increased over numbers in naïve mice in both sexes (**Figure 7C**). Effector immune cell infiltration, including total CD3^+^ T cells, CD4^+^ T cells, CD8^+^ T cells, and CD4^+^ regulatory T cells were increased over naïve levels in both sexes (**Figure 7D**). Remarkably, the only sex-specific difference with respect to immune cell infiltration was in the number of tumor antigen-specific CD8^+^ OT1 T cells, in which male mice had greater numbers of these cells in their bladder (**Figure 7D**).

**Figure 7.**
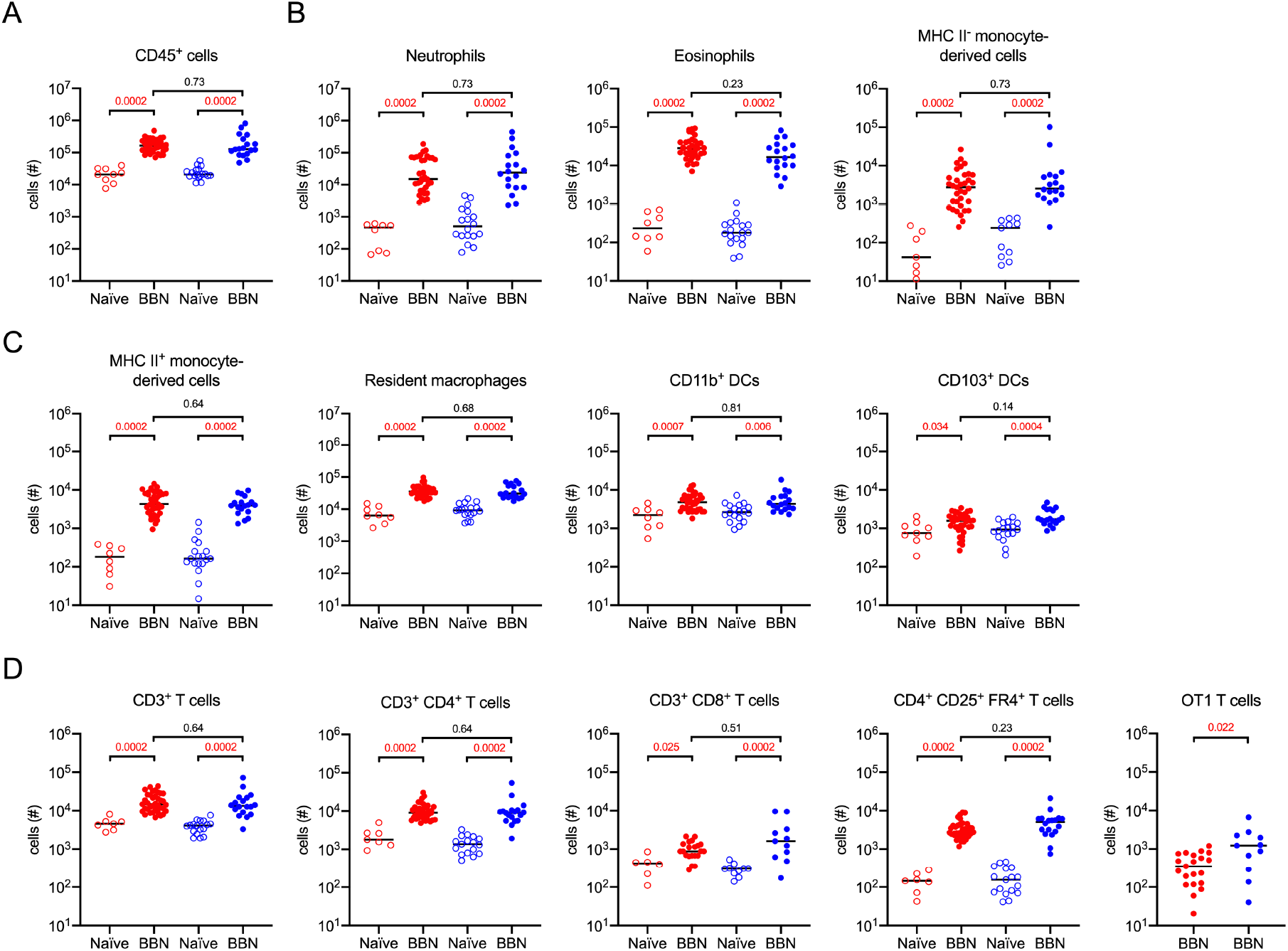
More CD8^+^ tumor-specific T cells infiltrate male bladders than female bladders. Mice were treated as shown in Figure 5A. Graphs depict the total number of specified immune cell populations in naïve or tumor-bearing female (red dots) and male (blue dots) URO-OVA mice. Graphs are pooled from 10 experiments, *n*=1-7 mice per experiment. Each dot represents 1 mouse, lines are medians. Significance was determined using the nonparametric Kruskal-Wallis test to compare naïve to BBN-treated mice for each immune cell population and to compare female to male BBN-treated mice. p-values were corrected for multiple testing using the FDR method. All calculated/corrected p-values are shown and those meeting the criteria for statistical significance (p<0.05) are depicted in red.

## Discussion

Our aim in this study was to develop optimized bladder cancer models in which tumor antigen-specific immunity can be studied during tumor development and treatment with known and novel therapeutic agents and combinations. After incorporation of the same model antigen, ovalbumin, into the MB49 orthotopic model and the BBN carcinogen model, we compared immune responses in these two different, but commonly used bladder cancer mouse models. These models are intended to help the bladder cancer research community to dissect how features of the tumor, including infiltrating immune cells and associated microenvironment, contribute to and are impacted by therapeutic approaches to support tumor immunity. With this knowledge, there is the potential to rationally design combination therapeutic approaches to reduce the incidence of BCG unresponsiveness or tumor recurrence, specifically in women and men. In sum, we found that using the URO-OVA mouse in the BBN carcinogenic model permitted the measurement of relevant cell surface molecules and immune cell infiltration in both female and male mice, leading to the identification of a potential sex-mediated difference in the infiltration of tumor-specific T cells.

We began by investigating the impact of BCG vaccination on the immune response to BCG therapy. Whether to include BCG vaccination as a part of bladder cancer patient treatment is debated. Vaccination was included in the original BCG therapy protocol because purified protein derivative (PPD) seroconversion, from PPD- to PPD+, correlated with positive clinical outcomes (Kelley et al., 1985; Winters and Lamm, 1981). BCG vaccination was abandoned, however, when it was demonstrated that percutaneous vaccination concurrent with intravesical therapy did not impart additional benefit to patients (Luftenegger et al, 1996). We found increased infiltration of monocyte-derived cells and CD4^+^ T cells when BCG-vaccinated nontumor-bearing mice were instilled with BCG three times, however CD8^+^ T cell infiltration was not different from that observed in the control PBS-injected, tumor-naïve mice instilled with BCG three times, suggesting that benefits from vaccination may be overcome by repeated instillation. BCG vaccination did not impact the infiltration of total CD4^+^ or CD8^+^ T cells into MB49 tumor-bearing mice treated with BCG three times, nor did it alter infiltration of any immune cell subset measured in the BBN tumor model. As BCG-vaccinated mice survive MB49 orthotopic bladder tumor challenge better than nonvaccinated mice when treated intravesically with BCG (Biot et al., 2012), our findings suggest that the mechanism of survival in this model is not a result of improved immune cell infiltration, but may be due to differences in the type or quality of the immune response induced.

It was disappointing that it was not possible to use male mice in the MB49 orthotopic model, as this is a commonly used model with tumors developing over weeks, rather than months as in the BBN model. We did recapitulate historical findings that BBN induces tumor growth at a faster rate in male mice compared to female mice (Bertram and Craig, 1972; Kaneko and Li, 2018) in the URO-OVA transgenic mouse, permitting use of this model to specifically explore sex differences in the immune response to immunotherapy in the BBN model. Greater understanding of the influence of sex and gender are needed in bladder cancer as men make up the majority of bladder cancer patients, although women are often diagnosed with more advanced disease and have poorer outcomes (Marks et al., 2016; Uhlig et al., 2018). Sex hormone receptors play a role in bladder cancer, as transgenic androgen receptor expression in the bladder increases tumor development in male and female mice, whereas mice deficient for the androgen receptor, specifically in the urothelium, are protected from BBN-induced tumorigenesis (Hsu et al., 2013; Johnson et al., 2016). In addition to sex hormone influence, higher expression of the X-linked gene KDM6A (lysine-specific demethylase 6A) is correlated with reduced risk of bladder cancer in female mice and women (Kaneko and Li, 2018).

Whether specific differences in the immune response to tumor growth or therapy exist between the sexes is largely unknown. We had anticipated finding differences in innate immune cell infiltration in the BBN model, as we reported that neutrophils, eosinophils, and monocytes infiltrate to a greater extent in female mice with bladder infection compared to infected male mice (Zychlinsky Scharff et al., 2019). While these cell types robustly infiltrated bladders of BBN-treated mice, the only sex-specific difference that we measured was a greater number of tumor antigen-specific CD8^+^ OT1 T cells in male mice compare to female mice. This observation may provide a first clue as to why men experience fewer tumor recurrences compared to women (Uhlig et al., 2018). Certainly, additional sex-based immune-mediated differences are likely involved, such as cytokine expression or T cell polarization, and the BBN URO-OVA model will be useful to explore these possibilities in future studies.

The MB49 orthotopic model and the BBN carcinogenesis model have been used to study bladder cancer for more than 50 years. Each has its strengths and weaknesses. Tumors in the MB49 model develop rapidly, however MB49 cell lines vary considerably from lab to lab and in some cases, display a rapid aggressive growth rate that precludes studies of adaptive immunity (El Behi et al., 2013; Saito et al., 2018). In addition, the tumor that develops is homogeneous likely not accurately reflecting human bladder cancers (Saito et al., 2018). The induction of bladder cancer by BBN leads to more heterogeneous tumors, which reflect human disease (Fantini et al., 2018; Sfakianos et al., 2020). However, tumors require months to develop and it is challenging to easily discern the stage and grade of tumors without sacrificing animals. With the introduction of the same antigen, the impact of different interventions on tumor-specific immunity can be directly compared between these models, potentially revealing additional important advantages or disadvantages in these models.

Given that NMIBC accounts for approximately 75% of bladder tumors, of which 30-50% will not respond to BCG therapy for unknow reasons; and that a BCG shortage has been in existence since 2012 (Messing, 2017), new therapies are urgently needed (Botteman et al., 2003; Fernandez-Gomez et al., 2009; Kamat et al., 2018; Witjes, 2006). Novel immunotherapeutic approaches, such as checkpoint blockade, that show considerable promise in muscle invasive bladder cancer are currently being tested to determine whether they will also positively impact NMIBC patient response, particularly in the subset of individuals who do not derive benefit from BCG therapy (Brahmer et al., 2012; Hamid et al., 2013; Powles et al., 2014; Rosenberg et al., 2016; Topalian et al., 2012). Indeed, improved immunity may only be possible with combination therapy, targeting different aspects of the immune response (*e.g.*, trafficking, priming) or the tumor *(e.g.*, stroma, infiltrating cells). For example, a-PD-1 antibody treatment slows tumor growth and extends survival compared to untreated or control isotype antibody treated mice in an MB49 subcutaneous tumor model, however, all mice except one developed tumors by the end of the experiment (Vandeveer et al., 2016). In this same study, a-PD-1 antibody treatment reduced bladder weight and *in vivo* radiance of MB49 luciferase-expressing tumors in mouse bladders, however, 50% of treated mice still died (Vandeveer et al., 2016). Interestingly, additional decreases in radiance and bladder weight were not observed when BCG was used in combination with a-PD-1 antibody treatment. Here, we did not observe an increase in overall T cell infiltration upon BCG therapy in BCG vaccinated MB49 tumor-bearing mice. However, we did observe a significant increase in CD8^+^ T cell infiltration, which is critical for antitumor activity, in vaccinated mice with combination BCG and α-PD-L1 therapy, suggesting a-PD-1 therapy and α-PD-L1 therapy may synergize differently.

Our intention is that these models be used by the bladder cancer research community to help establish a rational basis for therapeutic intervention, such as targeted depletion of specific immune cells or inhibition of immunosuppressive cytokines as alternative or combination treatments. Understanding how resident and recruited immune cell populations, such as macrophages, DCs, and T cells modulate response to therapy and identification of suppressive pathways will aid development of novel combination strategies to improve patient response. Moreover, these models will provide valuable information to understand sex-differences in bladder cancer immunity, as well as evidence to help development of new treatments for muscle invasive bladder cancer as an alternative to surgical resection and chemotherapy.

## Materials and Methods

### Ethics statement

Experiments were conducted at Institut Pasteur in accordance with approval of protocol number 2016–0010 by the *Comité d’éthique en expérimentation animale Paris Centre et Sud* (the ethics committee for animal experimentation), in application of the European Directive 2010/63 EU. Mice were anesthetized by injection of 100 mg/kg ketamine and 5 mg/kg xylazine and sacrificed by carbon dioxide inhalation. Groups were determined by partitioning of animals into cages upon their arrival to the animal facility.

### BCG substrain stocks

BCG Connaught, Tice, Danish, China, Japan, and RIVM were expanded from commercial preparations. For preparation of stocks used for vaccination and intravesical instillation, BCG strains were grown shaking (100 RPM) at 37°C in 7H9 Middlebrook medium (Sigma-Aldrich), complemented with ADC supplement (Sigma-Aldrich) and 0.2% glycerol (Sigma-Aldrich) for 10 days when BCG reached exponential growth phase. The BCG was pelleted, washed with PBS, and pelleted again. Pellets were resuspended in 5 ml PBS and BCG aggregates disassociated in GentleMACS M tubes using two cycles of the RNA01.01 setting on a GentleMACS machine (Miltenyi Biotech). The OD600 was determined, cultures diluted with PBS to 3×10^7^ colony forming units (CFU)/ml, and 1 mL stocks were aliquoted into cryovials (Nunc) and stored at −80°C until use. The number of viable BCG in stocks was determined by serial dilution and plating on 7H11 Middlebrook plates (Sigma-Aldrich), containing OADC supplement (Sigma-Aldrich) and 0.25% glycerol, at the time of each use (vaccination or intravesical instillation). Plates were incubated at 37°C for 2-4 weeks and colonies counted.

### Cell culture and MB49 transduction

Plat-E cells (Cell Biolabs), a retroviral packaging cell line (Morita et al., 2000), were thawed, cultured in D10 medium (DMEM (Life Technologies SAS), 10% fetal calf serum, 1mM HEPES buffer (Life Technologies SAS), 1mM sodium pyruvate (Life Technologies SAS), 1% 100X non-essential amino acids (Life Technologies SAS)) at 37°C, 5% CO_2_, for 2 days, then passaged 1:2 and cultured overnight in R10 medium (1X Roswell Park Memorial Institute (RPMI, (Life Technologies SAS), 10% fetal calf serum, 1mM HEPES buffer, 1mM sodium pyruvate, 1% 100X non-essential amino acids) plus 1μg/ml puromycin and 10μg/ml blasticidin (Invitrogen)). Plat-E cells were transfected with a plasmid encoding mCherry-ovalbumin fusion (Yatim et al., 2015) with Lipofectamine 2000 (Invitrogen) according to the manufacturer’s instructions, and incubated for 2 days. The culture supernatant was removed, centrifuged for 4 minutes at 1500 g, 4°C, and stored at −80°C. MB49 cells expressing luciferase (Summerhayes and Franks, 1979) were cultured in a 6-well plate in R10 medium for 3 days before transduction. For transduction, media was removed from the MB49 cells, the virus-containing supernatant was thawed, mixed with polybrene (Sigma-Aldrich) at a concentration of 0.1%, added to the MB49 cells, and incubated at 37°C, 5% CO_2_ for 1 hour. Following incubation, the plate was centrifuged at room temperature for 1½ hours at 1000 g, incubated for 2 hours at 37°C, 5% CO_2_, and then resuspended in R10 medium. mCherry positive cells were sorted using a BC FACSAria II cell sorter. mCherry expression was confirmed by flow cytometry. R10 medium was used to grow MB49 and MB49^mCh-ova^ cells. Cells were expanded and then frozen in bulk in 50% fetal calf serum, 10% DMSO, 40% R10 medium and stored in liquid nitrogen until use. Cells were allowed to recover at 37°C and 5% CO_2_ 2-4 days before tumor implantation.

### In vivo animal experiments

Mice: Male and female OT1 and URO-OVA transgenic mice (Barnden et al., 1998; Hogquist et al., 1994; Liu et al., 2007) were bred at the Institut Pasteur. URO-OVA mice were a kind gift from Yi Lou, University of Iowa. Male and female C57BL/6J mice were purchased from Charles River, France.

BCG vaccination: Animals were vaccinated subcutaneously at the base of the tail with 3×10^6^ colony forming units (CFU) of BCG Connaught in 100 μL PBS. Mock treated mice were injected with 100 μL PBS.

BCG-specific T cell determination: Spleens were passed through 70 μm filters (Miltenyi Biotech) into FACS buffer (PBS, 2% fetal calf serum, 0.2 mM EDTA), washed with PBS, and resuspended in 1X BD Pharm lyse buffer (BD Biosciences) for red blood cell lysis. Washed and pelleted splenocytes were resuspended in 1 mL FACS buffer, and 200 μL of splenocytes were mixed with fluorophore-conjugated BCG-specific tetramers (D^b^-restricted GAP (GAPINSATAM) peptide 100μg/sample, produced in house) and α-CD16/CD32 FcBlock (BD Bioscience) for 20 minutes. Cells were subsequently stained with antibodies to CD3, CD4, CD8 for 30 minutes. Samples were acquired on a BD LSRFortessa using DIVA software and data were analyzed by FlowJo software (Treestar).

Preparation of OT1 T cells: Spleens from transgenic OT1 transgenic mice were pressed through a 70 μm filter (Miltenyi Biotech), washed with PBS, and resuspended in 1X BD Pharm lyse buffer for red blood cell lysis. In the BBN model, at week 12, 10^6^ OT1 T cells in 100 μL of PBS were transferred to mice intravenously.

Orthotopic model: MB49 cell stocks were thawed and cultured for 2-4 days in R10 medium before implantation. For tumor implantation, mice were anaesthetized and bladders drained of urine by pressure to the lower abdomen. Bladders were pre-treated by transurethral instillation of 25 μL poly-L-lysine (0.1mg/ml in PBS (Sigma-Aldrich)) for 20 minutes. Poly-L-lysine was then aspirated and 2×10^5^ MB49 or MB49^mCh-ova^ cells (resuspended in 50 μL PBS) were instilled into the bladder. Cells were retained in the bladder for 20 minutes before catheter withdrawal.

Bioluminescence imaging: Both MB49 or MB49^mCh-ova^ cells express luciferase. In all mice instilled with tumors, tumor take was verified using the IVIS Spectrum In Vivo Imaging System on day 4 and day 7 post tumor instillation. The lower abdomen was shaved and animals were injected intraperitoneally (IP) with 150 μl of luciferin salt solution (33.3mg/ml) (Synchem) in PBS. After 5 minutes, mice were anesthetized by inhaled isoflurane and bioluminescence (total photons acquired) from tumor cells was visualized using the IVIS machine. Importantly, although treatments were begun on day 2 and/or day 3 post-tumor implantation, any animal that did not have a positive signal at day 7 post-tumor instillation was excluded from analysis.

Chemical induction model: Mice were given drinking water containing BBN (TCI America) at a concentration of 0.05%. BBN-containing water bottles (protected from light) were changed once per week until the end of the experiment.

Immunotherapy protocols: For therapeutic BCG intravesical instillation, MB49 tumor-bearing mice were anaesthetized at 2, 9, and 16 days post-MB49 implantation and BBN-treated mice were anaesthetized at weeks 12, 13, and 14 after the start of BBN administration. Their bladders were emptied and 5×10^6^ CFU of BCG Connaught was instilled intravesically in 50 μL PBS. Three instillations were used to avoid the necessity of sacrificing animals with obstructive tumors due to the rapid growth of MB49. Three instillations were maintained for the BBN-treated animals. For checkpoint inhibition, 200 μg of anti-PD-L1 antibody (clone 10F.9G2) were injected into the mouse IP. 24 hours after the last BCG treatment, mice were euthanized, and bladders collected, weighed, and analyzed by flow cytometry.

### Flow cytometry

Bladders were dissected, minced with scissors, and digested in 0.34 U/ml Liberase TM (0.06mg/ml, Roche) as previously described for 60 minutes at 37°C with manual agitation every 15 minutes (Zychlinsky Scharff et al., 2017). Cell suspensions were passed through a 100 μM filter (Miltenyi Biotech), and washed with PBS. Cell suspensions were stained with the Zombie Red fixable viability kit (Biolegend) prior to cell surface protein labeling according to the manufacturer’s instructions. Finally, cells were resuspended in FACS buffer (PBS, 2% fetal calf serum, 0.2 mM EDTA), incubated with 1% Fc block (CD16/CD32), and stained with the antibodies shown in **Table 1**. Cells were enumerated using AccuCheck counting beads (Invitrogen), according to the manufacturer’s instructions.

**Table 1:** Antibodies used for flow cytometry

| Antibody | Clone | Company |
| --- | --- | --- |
| **CD3** | 145-2C11 | BD Pharmingen |
| **CD3** | 500A2 | BD Pharmingen |
| **CD11c** | HL3 | BD Pharmingen |
| **CD25** | PC61 | BD Pharmingen |
| **Gr1 (Ly6C/Ly6G)** | RB6-8C5 | BD Pharmingen |
| **NK1.1** | PK136 | BD Pharmingen |
| **CD4** | RM4-5 | BD Horizon |
| **CD4** | GK1.5 | eBioscience |
| **CD8** | 53-6.7 | BD Horizon |
| **CD11b** | M1170 | BD Horizon |
| **CD45** | 30-F11 | BD Horizon |
| **CD45.1** | A20 | BD Pharmingen |
| **CD45.2** | 104 | BD Horizon |
| **CD103** | M290 | BD Horizon |
| **SiglecF** | E50-2440 | BD Horizon |
| **CD274** | MIH5 | BD OptiBuild |
| **CD279** | J43 | BD OptiBuild |
| **FR4** | eBio12A5 | eBioscience |
| **MHCII** | M5/114.15.2 | eBioscience |
| **F4/80** | MCA497A488 | Serotec/BIORAD |
| **CD64** | X54-5/7.1 | Biolegend |

### Statistical analysis

To plot graphs and determine statistical significance, GraphPad Prism V8 was used. The non-parametric Mann-Whitney test (2 groups) and Kruskal-Wallis test (3 or more groups) were used to determine statistical significance. In the case that more than 2 groups were being compared or to correct for comparisons made within an entire analysis or experiment, calculated p-values were corrected for multiple testing with the false discovery rate (FDR) method, (https://jboussier.shinyapps.io/MultipleTesting/), to determine the false discovery rate adjusted p-value. All calculated p-values are shown in the figures, and those that met the criteria for statistical significance (p<0.05) are denoted with red text.

## Author contributions

Conceptualization: MR, COB, EA, HZ, DC, MAI; Methodology: MR, COB, EA, HZ, DC, TC, AZS, MAI; Experimentation and data analysis: MR, COB, EA, HZ, DC, TC, AZS, MAI; Writing - Original Draft: COB, EA, HZ, DC, MAI; Writing - Review & Editing: MR, COB, EA, HZ, DC, TC, AZS, MAI; Funding Acquisition: MAI; Supervision: MR, MAI.

## Acknowledgments

We gratefully acknowledge insightful discussions, technical support, and/or critical reading of the manuscript by Dr John Taylor and Livia Lacerda Mariano. We thank our spouses and partners for being supportive of our work-at-home efforts during the COVID-19 pandemic. Funding: MAI was supported by funding from the The Leo & Anne Albert Institute for Bladder Cancer Care and Research, LabEx ImmunoOnco (ANR 10-LABX-0015), and a Projets Fondation ARC award. We also thank the Erasmus+ programme and the University of Glasgow, and in particular, Drs Simon Milling and Robert Nibbs, for the academic opportunity offered through the integrated Masters (MSci) program with work placement in Life Sciences, which supported the research visits of COB, EA, and HZ.

